# Transcription factor dynamics reveals a circadian code for fat cell differentiation

**DOI:** 10.1101/245332

**Authors:** Zahra Bahrami-Nejad, Michael L. Zhao, Stefan Tholen, Devon Hunerdosse, Karen E. Tkach, Sabine van Schie, Mingyu Chung, Mary N. Teruel

**Affiliations:** Department of Chemical and Systems Biology, Stanford University, Stanford, CA, 94305, USA.

**Keywords:** Positive feedback, adipogenesis, circadian filtering, hormone oscillations, glucocorticoids, PPARG, CEBPB

## Abstract

Glucocorticoid and other adipogenic hormones are secreted in mammals in circadian oscillations. Loss of this circadian oscillation pattern during stress and disease correlates with increased fat mass and obesity in humans, raising the intriguing question of how hormone secretion dynamics affect the process of adipocyte differentiation. By using live, single-cell imaging of the key adipogenic transcription factors CEBPB and PPARG, endogenously tagged with fluorescent proteins, we show that pulsatile circadian hormone stimuli are rejected by the adipocyte differentiation control system, leading to very low adipocyte differentiation rates. In striking contrast, equally strong persistent signals trigger maximal differentiation. We identify the mechanism of how hormone oscillations are filtered as a combination of slow and fast positive feedback centered on PPARG. Furthermore, we confirm in mice that flattening of daily glucocorticoid oscillations significantly increases the mass of subcutaneous and visceral fat pads. Together, our study provides a molecular mechanism for why stress, Cushing’s disease, and other conditions for which glucocorticoid secretion loses its pulsatility can lead to obesity. Given the ubiquitous nature of oscillating hormone secretion in mammals, the filtering mechanism we uncovered may represent a general temporal control principle for differentiation.

**HIGHLIGHT:**

- We found that the fraction of differentiated cells is controlled by rhythmic and pulsatile hormone stimulus patterns.
- Twelve hours is the cutoff point for daily hormone pulse durations below which cells fail to differentiate, arguing for a circadian code for hormone-induced cell differentiation.
- In addition to fast positive feedback such as between PPARG and CEBPA, the adipogenic transcriptional architecture requires added parallel slow positive feedback to mediate temporal filtering of circadian oscillatory inputs

## INTRODUCTION

Slow, ongoing terminal cell differentiation is essential for replacing aging or damaged cells and for maintaining tissue size in all adult mammals. For example, adipocytes (fat cells) and cardiomyocytes renew in humans at rates of approximately 8% and 1% per year, respectively (Bergmann et al., 2009; Spalding et al., 2008). Since terminally-differentiating cells are often derived from large pools of precursor cells, differentiation is expected to be a rare event. In the case of fat cell differentiation, or adipogenesis, for which there are estimates of about one preadipocyte for every five differentiated cells (Tchoukalova et al., 2004), fewer than 1% of preadipocytes are believed to embark on a differentiation path on any given day under normal, homeostatic conditions. How such low rates of differentiation can be reliably maintained is puzzling given that preadipocytes are subjected to daily high increases of differentiation-inducing hormones such as glucocorticoids which are needed in mammals to mobilize energy and increase physical activity, but which also have been shown to strongly accelerate adipogenesis in vitro and in vivo (Campbell et al., 2011; Farmer, 2006; Lee et al., 2014; Park and Ge, 2017).

Glucocorticoids are secreted in healthy mammals in daily oscillatory patterns (Figure 1A)(Weitzman et al., 1971), as well as in sporadic bursts in response to stress. Levels of other adipogenic hormones such as ghrelin and prolactin, that raise cAMP, and insulin, fluctuate in vivo as well. Given that so few preadipocytes differentiate each day despite significant daily increases in adipogenic hormone stimuli, we hypothesized that the regulatory circuit controlling adipocyte differentiation might be filtering out circadian and short hormone stimuli and that differentiation would only start to occur if the duration of the trough between circadian pulses shortens or if the hormone signal remain continuously elevated. Such a temporal filtering of hormone signals is consistent with the observation that, in contrast to short or daily oscillatory signals, more continuous glucocorticoid signals have been linked to increased fat mass (Campbell et al., 2011; Dallman et al., 2000; Lee et al., 2014). Shortening of the trough between glucocorticoid pulses, or “flattening of the daily oscillations” (Leliavski et al., 2015; Windle et al., 2013), due to irregular feeding or sleep cycles, prolonged treatment with glucocorticoid hormones, chronic stress, or metabolic diseases such as Cushing’s disease have all been shown to closely correlate with increased obesity (Balbo et al., 2010; Dallman et al., 2000; Lee et al., 2014; Panda, 2016).

**Figure 1.**
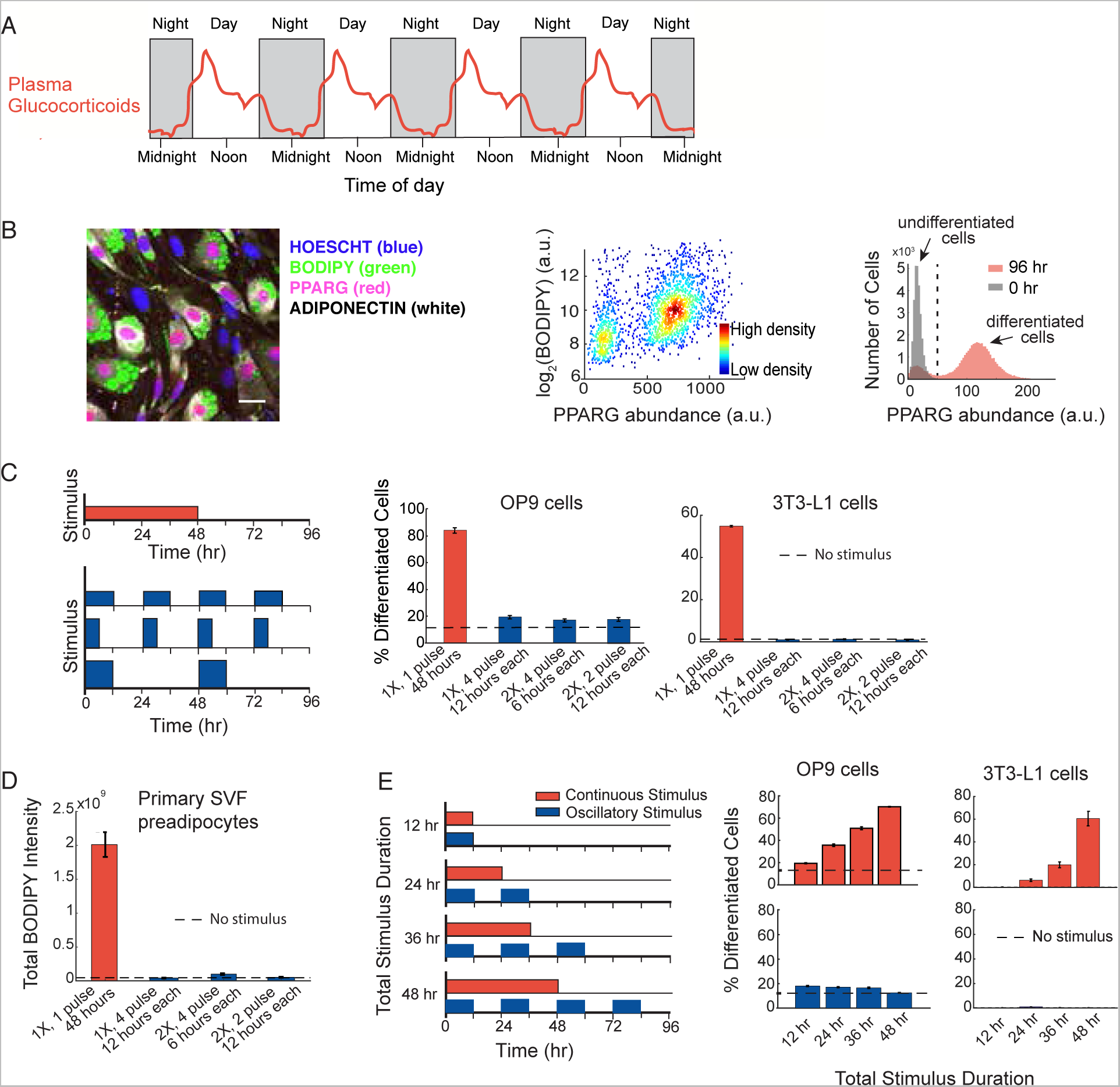
Preadipocytes reject circadian and rhythmic glucocorticoid stimuli. (A) Schematic depicting averaged glucocorticoid time courses in humans (adapted from Weitzman et
al, 1971). (B) Single-cell immunofluorescence assay used to quantitate percent of differentiated cells based on Park et al, 2012. Preadipocyte cells were fixed and stained with Hoechst to visualize nuclei (blue), BODIPY to visualize lipids (green), and antibodies against PPARG (red) and adiponectin (white). The latter 3 signals are closely correlated in individual cells. Images show mouse OP9 cells. Scale bar, 10μm. Scatter plot shows that high-PPARG cells display mature fat cell features and accumulate high levels of lipid. The image and histograms show that application of an adipogenic stimulus causes cells to split into two populations of low PPARG and high PPARG cells. The middle of the trough between the bimodal PPARG intensity peaks (shown by black dotted line) was used to separate undifferentiated and differentiated cells. Percent of differentiated cells was assessed at t = 96 hours after applying adipogenic stimulus, using approximately 7,000 cells per technical replicate. See also Figure S1. (C) Different patterns of DMI stimuli were applied to OP9 and 3T3-L1 preadipocyte cells: 48 hours continuous delivery (red) and 3 different pulsatile protocols (blue): 12 hours on/12 hours off, 6 hours on/18 hours off, and 12 hours on/36 hours off. The concentration of DMI stimulus used for a 48-hour continuous pulse was 1μM dexamethasone (dex), 250 μM IBMX, and 1.75 nM insulin. To keep the total amount of stimuli, i.e. area under the curve, constant for all 4 protocols over the 96-hour experimental timeframe, the concentration of dex and IBMX was increased proportionally to compensate for decreases in pulse duration. To end a pulse, the DMI containing media was aspirated, cells were washed gently three times with fresh media, and the cells were left in media with no glucocorticoids or IBMX, but containing 1.76 nM insulin. The doses of dex used were not saturating (Figures S2A and S2B), and using corticosterone instead of dex in the stimulus showed the same filtering effects (Figure S2C). (D) Experiments in primary SVF preadipocytes show that the same filtering effects are also observed in primary cells. SVF preadipocytes were plated into 96-well wells and induced to differentiate using the same pulsing protocols from (C). The Total BODIPY Intensity in each well was measured by imaging. (E) Applying continuous stimuli with durations greater than 12 hours resulted in corresponding increases in the number of differentiated OP9 and 3T3-L1 cells. However, the same total stimuli given in circadian pulses resulted in only minimal differentiation. (C-E) Barplots represents mean +/- s.e.m. from 3 technical replicates. All data shown are representative of 5 independent experiments.

Adipogenesis occurs over several days by activation of a core adipogenic transcriptional network that is similar in vivo and in vitro (Roh et al., 2017; Rosen and Spiegelman, 2014). A critical feature of this hormone-induced differentiation process is its reliance on initiation processes requiring CEBPB and a canonical core positive feedback between PPARG and CEBPA that works together with additional secondary positive feedbacks and regulatory mechanisms (Ahrends et al., 2014; Farmer, 2006; Lefterova et al., 2014; Rosen and Spiegelman, 2014). The positive feedbacks are proposed to control a bistable switch that separates a distinct undifferentiated precursor state and a distinct differentiated state (Ahrends et al., 2014; Jukam and Desplan, 2010; Park et al., 2012; Wang et al., 2009). Nevertheless, while these previous studies provided indirect evidence for bistability using different strengths of continuous stimuli, they did not address the physiologically more relevant question if and how oscillating stimuli of the same total dose control the differentiation process. Futhermore, since these previous studies did not use live cell analysis, they were only able to infer a threshold and bistability of the differentiation process and could not directly show it. Since transcriptional processes are typically variable between cells, investigation of the dynamic processes controlling the switch requires time course-analysis in single cells (Loewer and Lahav, 2011; Spencer et al., 2009).

Here we investigated whether a differentiation system can temporally filter out circadian hormone inputs, a question that to our knowledge has not been addressed by previous studies which typically administered only constant levels of hormone stimuli or single hormone pulses. To test for dynamic control of differentiation, we employed a multi-day, live-cell imaging approach in which we monitored the expression levels of the transcription factors CEBPB and PPARG by endogenously tagging the respective loci in model mouse adipocyte precursor cells (OP9 cells) with fluorescent protein. Strikingly, we demonstrate a temporal control principle whereby preadipocytes reject normal pulsatile, daily hormone inputs due to a combined slow and fast positive feedback circuit that controls the self-amplification of PPARG. The fraction of cells that differentiate only starts to increase when the durations of hormone pulses become extended beyond normal circadian pulse durations.

## RESULTS

### Circadian and rhythmic hormone stimuli are rejected by the preadipocyte differentiation system

To determine whether and how pulsatile versus continuous hormone stimulation regulate adipogenesis, we used 3T3-L1 and OP9 *in vitro* models for adipocyte differentiation (Mandrup and Lane, 1997; Wolins et al., 2006), as well as primary stromal vascular fraction (SVF) preadipocytes isolated from mice (Luo et al., 2017). To induce differentiation, we applied pulses of an adipogenic hormone cocktail (DMI, see Methods) that mimics glucocorticoids and GPCR-signals that raise cAMP, such as ghrelin and prolactin. These signals have been shown to increase and decrease in circadian or fluctuating patterns in vivo (Carré and Binart, 2014; Thompson et al., 2004) and to promote adipogenesis by stimulating a core adipogenic transcriptional network that is similar in vivo and in vitro (Roh et al., 2017; Rosen and Spiegelman, 2014). Here we used the commonly-used DMI cocktail as a maximal stimulus of glucocorticoids and synergistic hormone signals.

We quantified differentiation by measuring expression of PPARG, the master transcriptional regulator of adipogenesis, whose expression in individual cells correlates closely with lipid accumulation, as well as with expression of GLUT4, adiponectin, and other mature fat cell markers (Cristancho and Lazar, 2011; Tontonoz and Spiegelman, 2008) (Figures 1B, S1A-S1C). To determine the effect of pulsatile versus continuous DMI stimulation on adipogenesis, we used a protocol in which we added and removed DMI in four different pulse patterns while keeping the integrated DMI stimulus constant over four days (Figure 1C, left). Strikingly, rhythmic cycles of DMI stimuli applied to preadipocytes with durations of 12 hours or less caused only minimal differentiation (Figure 1C, blue) while, in contrast, the same total amount of stimulus applied continuously caused robust differentiation (Figure 1C, red; dashed line reflects basal differentiation). To validate that this is a general phenomenon, we applied the same protocols to primary SVF preadipocytes isolated from mice and measured accumulation of lipids which is a commonly used as a marker for differentiation in these cells (Rodeheffer et al., 2008; Van et al., 1976). Markedly, the primary preadipocytes showed an equally strong failure to differentiate in response to daily 12-hour on/12-hour off DMI stimulation compared to robustly differentiating when the same total integrated DMI stimuli was applied in a continuous manner for 48 hours (Figures 1D and S1D).

Increasing the continuous stimulus durations from 12 to 24, 36, and 48 hours resulted in a progressive increase in differentiated cells (Figure 1C, red). However, once again, when the same total amount of stimulation was applied in circadian cycles, minimal differentiation was observed in OP9 and 3T3-L1 cells (Figure 1C, blue), as well as in primary SVF preadipocytes (Figure S1E). Control experiments confirmed that the doses of applied DMI stimuli were not saturating and that even the application of much stronger doses of DMI failed to induce differentiation when applied in circadian and rhythmic pulses (Figures S2A and S2B). The same filtering and differentiation response to pulsatile versus continuous stimuli was observed whether a synthetic glucocorticoid, dexamethasone, or the physiological glucocorticoid, corticosterone, was used in the hormone stimulus cocktail (Figure S1C). Taken together, these data show that preadipocytes make use of a robust filtering mechanism to maintain low rates of differentiation when exposed to daily hormone pulses. Intriguingly, the near complete filtering of differentiation-inducing stimuli was observed for pulse durations of up to 12 hours (Figure 1E), approximately reflecting the duration of normal circadian glucocorticoid patterns that preadipocytes are expected to experience in vivo (Weitzman et al., 1971) (Figure 1A).

### CEBPB mirrors the dynamic changes in pulsatile hormone inputs

To understand at which step in the differentiation signaling pathway input signals are filtered out, we compared the effect of two different stimuli: (i) rosiglitazone which binds to and directly activates PPARG (Lehmann et al., 1995) and (ii) DMI which mimics glucocorticoid stimuli and increases transcription of PPARG (schematic shown in Figure 2A, (Chawla et al., 1994)). The stimuli were applied for different durations ranging from 2 to 48 hours. When PPARG was directly activated with rosiglitazone, there was no filtering of input signals, and the fraction of differentiated cells increased proportionally with increasing pulse durations (Figure 2B, blue bars). However, when PPARG was indirectly activated with DMI, preadipocytes showed an increase in differentiation only for pulses longer than 12 hours (Figures 2B and S2D, red bars), arguing that the observed filtering of glucocorticoid input stimuli occurred before or along with PPARG activation.

**Figure 2.**
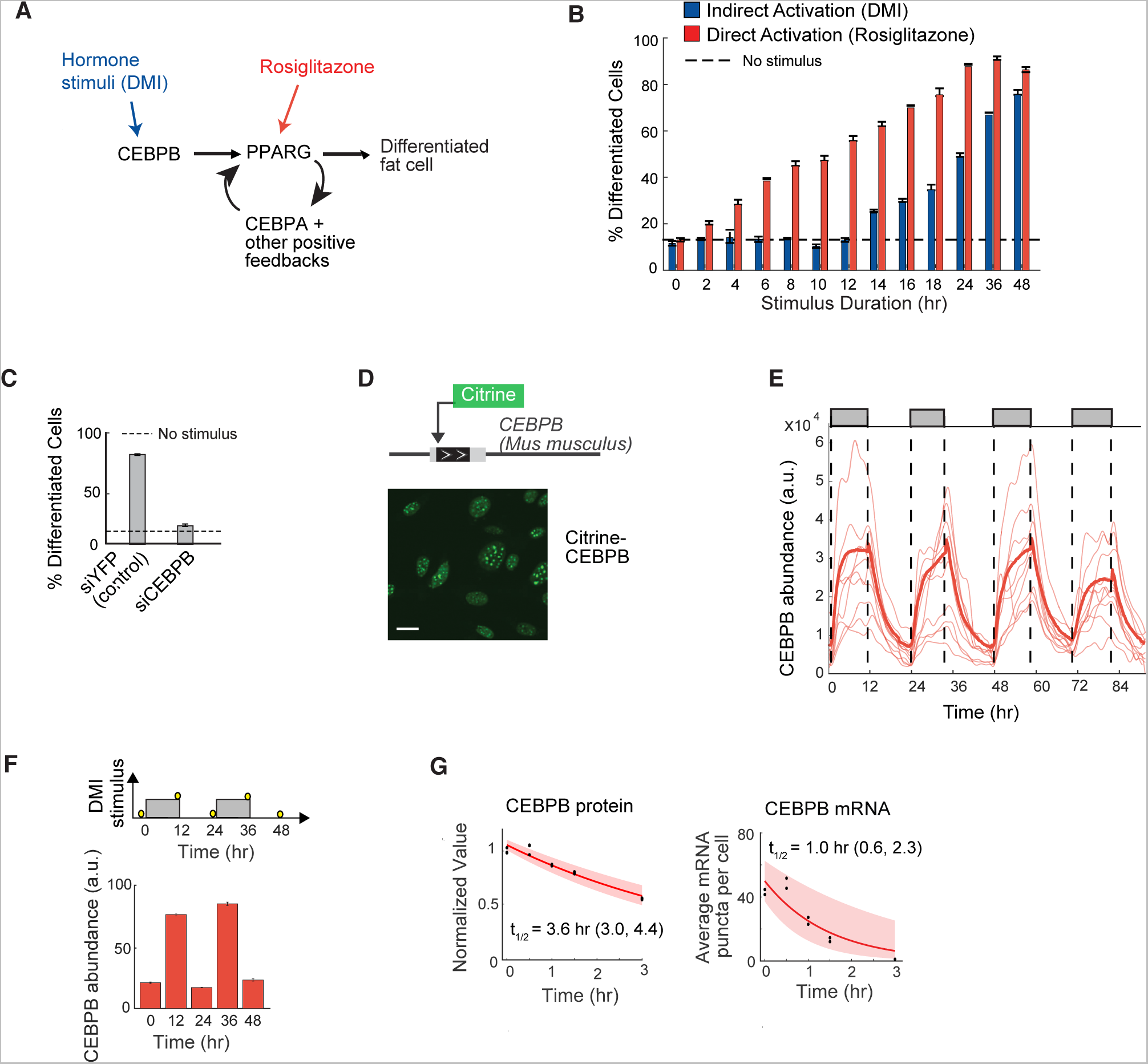
CEBPB mirrors the dynamics of the hormone input and transmits the pulse durations to PPARG. (A) Simplified schematic of the canonical adipogenic transcriptional network (Farmer, 2006; Rosen and Spiegelman, 2014). (B) Stimuli pulses of different durations were applied to OP9 cells. DMI was used to indirectly activate PPARG, and rosiglitazone was used to directly activate PPARG. Data from 3T3-L1 cells is shown in Figure S2D. (C) Suppression of differentiation by siRNA-mediated depletion of CEBPB versus control (YFP) in OP9 cells. (B, C) The percent of differentiated cells was measured as in Figure 1B and show mean +/- s.e.m. from 3 technical replicates, representative of 3 independent experiments. (D) Endogenous CEBPB (LAP* isoform) in OP9 cells was tagged with citrine (YFP) using CRISPRmediated genome editing. Scale bar, 10 μm. (E) Single-cell time courses of total citrine-CEBPB nuclear abundance (CEBPB abundance) in response to a 12-hour on / 12-hour off pattern of DMI stimulus (thin lines). The bold line represents the population median and shaded regions represents the 25th to 75th percentile of the population, n > 300 cells. Citrine-CEBPB expression increased with t_1/2_ ∼ 2 hours with stimulus addition and decreased with t_1/2_ ∼3 hours with stimulus removal, as estimated by a single exponential decay model. (F) 3T3-L1 cells were stimulated with circadian pulses (12-hour on / 12-hour off) of DMI and fixed at the respective time points marked with yellow circles (top). Cells were stained with anti-CEBPB antibody and Hoescht as a nuclear marker and imaged. The mean nuclear CEBPB abundance was quantitied in each cell by measuring the intensity of the anti-CEBPB antibody. Each bar represents the mean +/- s.e.m. of 3 technical replicates, each with approximately 500 cells. The barplot shown is representative of 3 independent experiments. (G) OP9 cells were stimulated with DMI for 24 hours, treated with cyclohexamide or actinomycin-D, fixed at the plotted time points, and immunofluorescence or RNA FISH was performed to measure CEBPB protein levels or count the number of mRNA puncta per cell, respectively. First-order exponential decay curves were fitted to data (t_1/2_ ∼3.6 hr and 1 hr, respectively). The shaded region represents the 95th confidence bounds.

Since the transcription factor CEBPB is induced by DMI stimuli and is necessary for PPARG expression (Yeh et al., 1995) (Figures 2A and 2C), we tested whether CEBPB mediates the filtering of hormone pulses. We measured dynamic changes in CEBPB levels by using CRISPR-mediated genome editing to generate an OP9 preadipocyte cell line with Citrine (YFP) fused to the N-terminus of endogenous CEBPB (Figures 2D and S3). Control experiments verified that the tagged endogenous protein was regulated similarly and had the similar expression levels and lifetimes to the untagged endogenous CEBPB protein (Figure S5). To obtain timecourses of CEBPB levels, cells were automatically tracked over several days, and their nuclear YFP fluorescence was recorded. Surprisingly, these live-cell imaging experiments showed that the nuclear abundance of CEBPB was highly dynamic and closely mirrored the hormone input for oscillating stimuli (Figure 2E). Nuclear abundance of CEBPB has been shown to reflect DNA binding of CEBPB (Siersbæk et al., 2011), arguing that not only the level but also the activity of CEBPB follows the stimulus dynamics. Similar nuclear CEBPB dynamics as in Figure 2E were observed in 3T3-L1 cells stimulated with a pulsatile pattern (Figure 2F), suggesting that fast CEBPB dynamics mirror the external hormone stimulus in different pre-adipocyte models. The observed fast changes in CEBPB levels can be explained by the fast degradation rates of both CEBPB protein and mRNA (Figures 2G and S5).

### Live-cell imaging of fluorescently-tagged endogenous PPARG reveals the existence of a threshold where internal positive feedback becomes independent of external stimuli

Because CEBPB dynamics closely mirrored the hormone input stimuli, filtering must occur downstream of CEBPB, suggesting that PPARG might be involved. We again made use of CRISPR-mediated genome editing to generate an OP9 cell line with citrine (YFP) fused to the N-terminus of endogenous PPARG (Figures 3A and S3). Control experiments verified that the tagged endogenous protein was regulated similarly and had the similar expression levels and lifetimes to the untagged endogenous PPARG protein (Figure S6).

The role of PPARG in filtering circadian inputs has to be investigated in the context of previous work that showed that PPARG is a critical part of a bistable switch that converts preadipocytes to adipocytes (Park et al., 2012). Bistable switches between two fates are a fundamental part of differentiation processes, and many models predict that a threshold in the expression of a core regulator must exist that decides whether or not a cell will differentiate (Jukam and Desplan, 2010; Kalmar et al., 2009; Wang et al., 2009). However, the existence of a threshold has never been directly shown since to do so requires live-cell measurement of the presumed regulator in individual cells while being able to remove the differentiation-inducing stimulus and continuing to track individual cells to their final differentiation state. Such live-cell analysis is necessary in order to verify that reaching the threshold level of the presumed regulator indeed results in that cell eventually transitioning irreversibly into the differentiated state.

Our citrine-PPARG cells allowed us now to directly test for the first time whether such a threshold exists in cell differentiation. We continuously imaged and automatically tracked citrine-PPARG preadipocytes over a 4-day time course using a 48-hour continuous DMI stimulation protocol. Markedly, when the DMI stimulus was removed, the population of preadipocytes split into two distinct populations: cells in which PPARG levels caught on and continued to increase with cells reaching the differentiated adipocyte state (red), and cells in the same population in which PPARG level dropped back down with cells staying in the undifferentiated state (blue) (Figure 3B). As shown in Figure 2A, PPARG levels are regulated by both external hormone stimulus and cell-intrinsic positive feedback. This dual regulation of PPARG levels can be more clearly seen in the live-cell traces when cells are binned together according to their level of PPARG at the time of DMI removal (Figure 3C). Markedly, in cells whose PPARG level is above a critical threshold when DMI is removed (red cells), PPARG levels first fall partially down due to the loss of the external hormone stimulus and then increase again as the cell-intrinsic positive feedbacks engage to continue to raise PPARG levels, independently of external stimuli, to the fully differentiated state. In cells whose PPARG level does not increase to this critical threshold (blue cells), the cell-intrinsic positive feedbacks do not engage strongly enough and cells thus fall back to the undifferentiated state. Figure 3D shows measurements extracted from individual cell timecourses which show the PPARG level in each cell at 48 hours before DMI removal and then again in the same cell at 96 hours when the final differentiation state is known. The threshold can then be calculated as the midpoint between the two distributions of the PPARG level for differentiated and undifferentiated cells before DMI removal (Figure 3D). This calculated threshold (marked as a yellow triangle in Figures 3B-D) predicts differentiation outcome at 96 hours already at the time of DMI removal at 48 hours.

**Figure 3.**
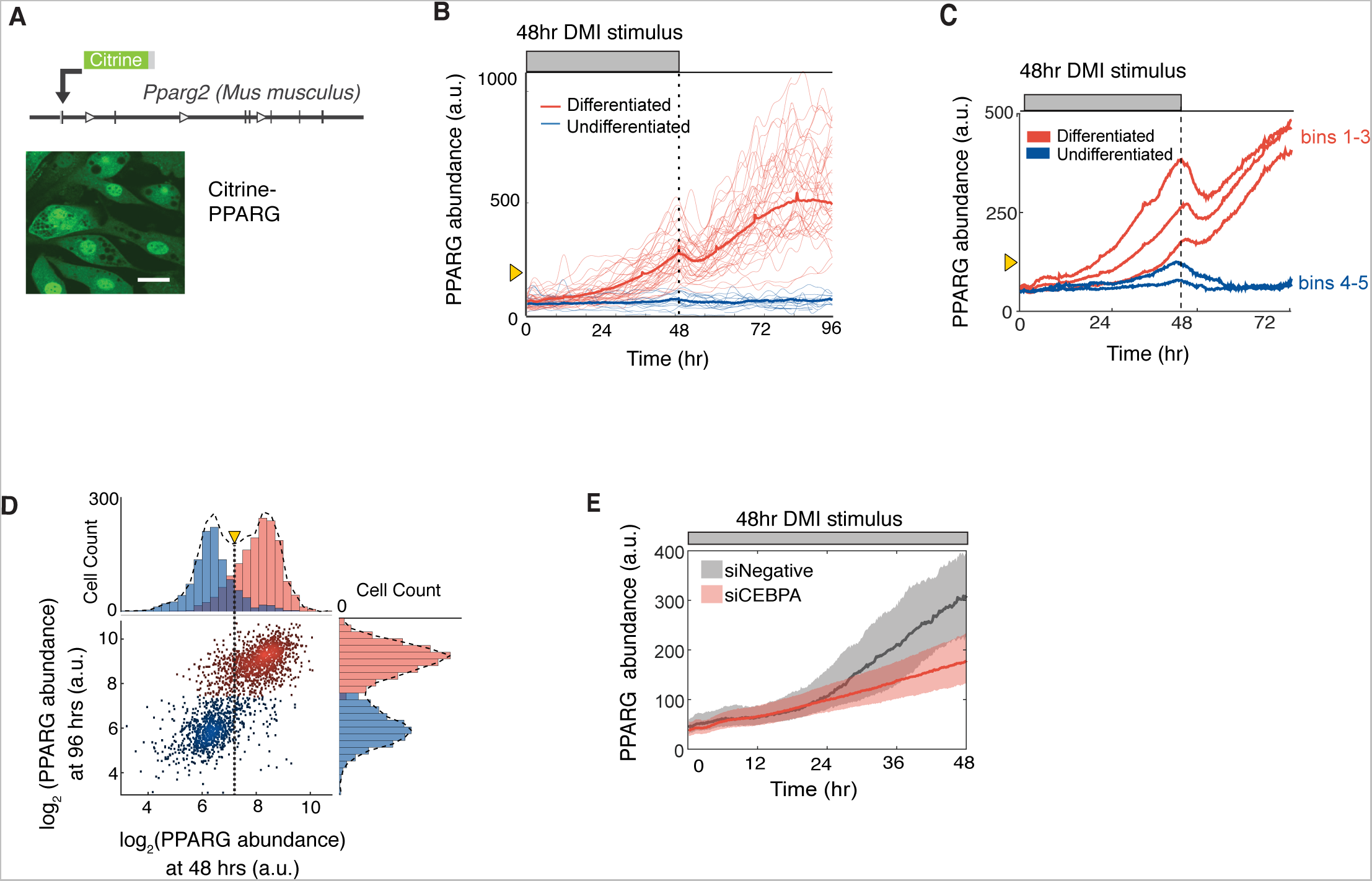
PPARG levels increase slowly and typically do not reach the threshold to differentiate unless hormone stimuli are applied for longer than 24 hours. (A) Endogenous PPARG2 in OP9 cells was tagged with citrine (YFP) using CRISPR-mediated
genome editing. Scale bar, 5 μm. (B) Identifying a PPARG threshold for differentiation by carrying out live-cell imaging of citrine-PPARG cells induced to differentiate with a DMI stimulation for 48 hours followed by insulin for another 48 hours. When the DMI stimulus is removed at 48 hours, cells either sustain or lose the increase in nuclear PPARG intensity (PPARG abundance). The thin lines show 50 representative single cell traces, and the bold line shows the population mean of 500 cells. Cells that differentiated after 96 hours are marked in red, and cells that stayed in the preadipocyte state are marked in blue. The yellow triangle shows the threshold calculated in Figure 3D. (C) Timecourses from 419 cells were divided into 5 groups (bins) according to their PPARG level at 48 hours, and the average of each bin was plotted. An initial drop was seen in each averaged timecourse after DMI removal, resulting in a return to the basal state for cells that had lower levels of PPARG and a delayed increase to the maximal state in cells that had higher levels of PPARG. (D) Scatter plot comparing the level of PPARG in the same cell measured at 48 hours just before DMI is removed and at 96 hours when PPARG levels reach their plateau in differentiated cells. The color of the dots and histograms mark cells that will be differentiated (red) or not differentiated (blue) at 96 hours, respectively. The histograms on top shows the existence of a clear PPARG threshold already at 48 hours before the stimulus is removed (marked with dotted line) that can distinguish between cells that will go on to differentiate from those that will not. The yellow triangle shows the threshold. (E) Live-cell imaging of citrine-PPARG cells following transfection with CEBPA-targeted or control(YFP) siRNA shows that knockdown of CEBPA, a main feedback partner of PPARG, results in loss of the feedback engagement point. Cells were stimulated to differentiate by adding DMI. Plotted lines are population median traces with shaded regions representing 25th and 75th percentiles of approximately 700 cells per condition.

We also observed in the analysis in Figure 3C that PPARG levels start to increase faster approximately 24 hours after addition of the DMI stimulus and long before the threshold for differentiation is reached. When we knocked down CEBPA, the core positive feedback partner of PPARG, this faster increase in PPARG was absent and cells failed to reach the threshold for differentiation (Figure 3E), arguing that the external hormone stimulus first gradually increases PPARG levels for about 24 hours before cell-intrinsic positive feedback to PPARG engages to then allow cells to reach the threshold for differentiation. The threshold level, which is reached after approximately 36-48 hours of continuous external hormone stimulation, is the level of PPARG at which cell-intrinsic positive feedback, initially triggered around 24 hours, has becomes sufficiently strong to sustain the increase in PPARG even in the absence of external hormone stimuli.

### The core adipogenic transcription factors PPARG and CEBPA rapidly degrade which keeps PPARG levels below the threshold to differentiate for daily oscillatory stimuli

Given the existence of a PPARG threshold, it was plausible that daily hormone stimuli fail to trigger differentiation due to a failure of PPARG levels to reach the threshold. Such a mechanism is plausible due to the very slow PPARG increase we observed in Figures 3B and 3C. It takes PPARG approximately 24-48 hours after continuous stimulation to reach the irreversible threshold for differentiation, meaning that shorter stimuli pulses may fail to trigger differentiation. Indeed, live-cell imaging experiments showed that repetitive 12-hour on / 12-hour off stimuli resulted in only minimal increases in PPARG levels in most cells (Figures 4A and 4B), and thus prevented differentiation compared to when an equally-strong continuous stimulus was applied (Figure 3B). Interestingly, when the stimulus was off during pulsatile condition. PPARG levels not only stopped to increase but also dropped (blue traces in Figures 4A and 4C). The drops between pulses prevent PPARG levels from gradually increasing to the threshold to trigger differentiation for repetitive pulse protocols. The rapid changes in PPARG levels can be explained by the short half-lives of PPARG protein and mRNA, approximately 1 and 0.8 hour respectively (Figure 4D).

**Figure 4.**
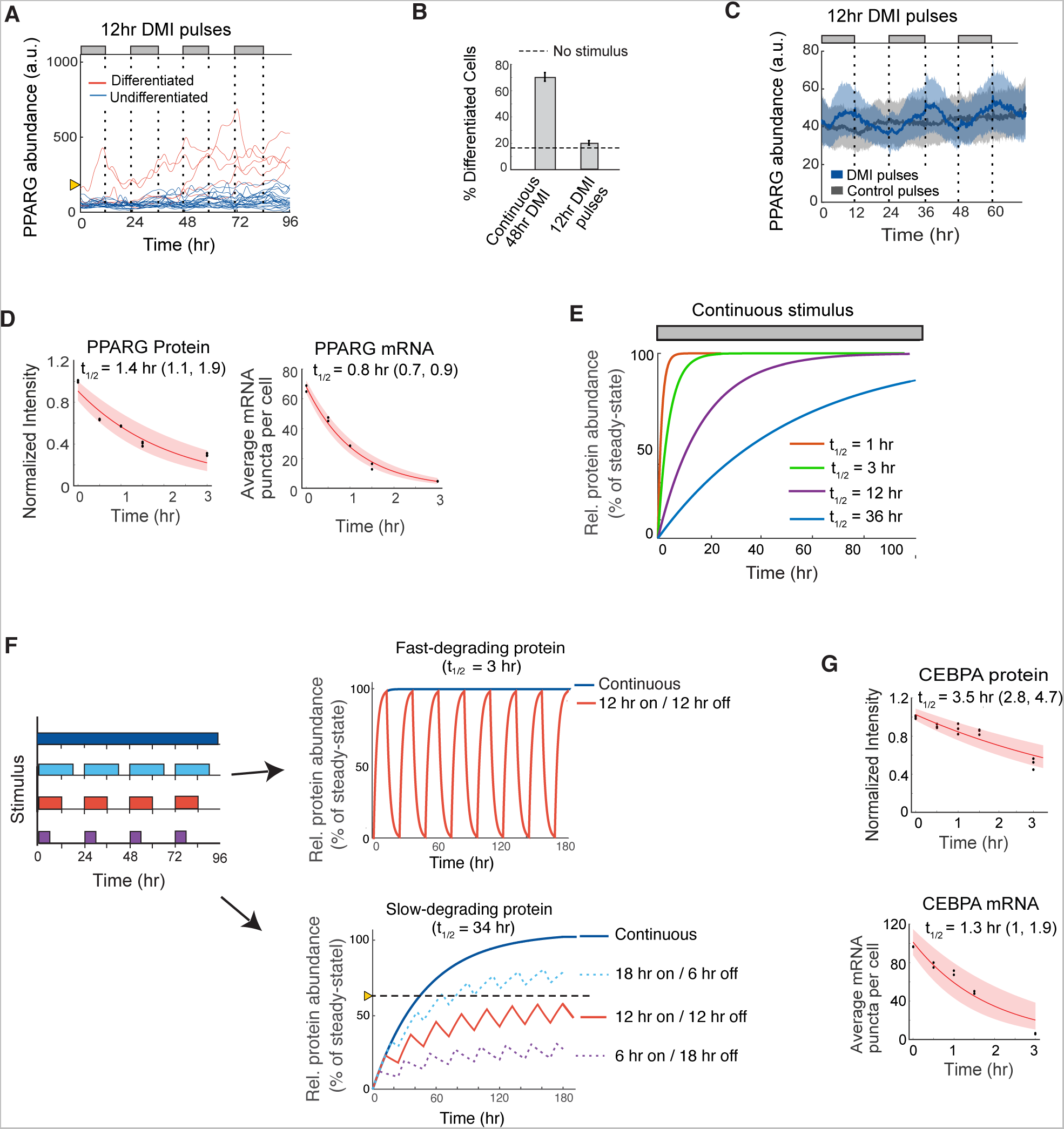
The core adipogenic transcription transcription factors PPARG and CEBPA rapidly degrade which keeps PPARG below the threshold to differentiate in response to oscillatory circadian hormone inputs. (A) Single-cell timecourses showing the dynamics of citrine-PPARG expression when adipogenic DMI stimuli are applied in a pulsatile 12-hour on/12-hour off manner over 4 days. Cells were marked as differentiated (red) and undifferentiated (blue) based on whether their final citrine-PPARG values was above or below the threshold for differentiation defined in Figure 3. (B) Comparison of the percent of differentiated cells resulting from continuous (Figure 3B) versus circadian (Figure 4A) stimuli. Error bars show mean +/- s.e.m. from 6 technical replicates, representing 3 independent experiments. (C) Zoom-in of the undifferentiated cells (blue traces) in (A). If PPARG levels do not reach the threshold to differentiate, PPARG levels drop back down to precursor levels each time the glucocorticoid and cAMP-inducing (DMI) stimulus is removed. (D) Lifetime analysis of PPARG protein and mRNA as in Figure 2G. (E) Plot of the times required to reach steady state for molecules with different half-lives and first order decay. (F) A slow-degrading regulator converts input stimuli with increasing pulse durations into increasing output signal amplitudes whereas a fast-degrading regulator outputs only a single maximal amplitude. Model output in response to different input pulse protocols shown on the left and assuming exponential increase and decay of proteins with t1/2 = 3 and 34 hrs, respectively. (G) Lifetime analysis of CEBPA protein and mRNA as in Figure 2G.

The fast degradation times of CEBPB and PPARG and the failure of PPARG levels to sufficiently increase under pulsatile conditions can explain the filtering seen in response to daily hormone pulses. However, the fast dynamics also raised the puzzling question of how PPARG levels can slowly build over days to result in differentiation when cells are subjected to continuous adipogenic stimuli. It has previously been shown that 2 days of stimulation are needed to raise PPARG levels to be able to convert a large fraction of preadipocytes to the point where they stay on a differentiation path even when the stimulus is removed (Chawla et al., 1994; Tontonoz et al., 1994). As shown in Figure 3B, it takes most cells 24-48 hours to build PPARG levels to the threshold to differentiate. Since the time for a protein to accumulate to a steady-state level depends only on its degradation rate when the synthesis rate is not changing (Rosenfeld et al., 2002), a regulatory circuit built from only fast-degrading proteins such as PPARG and CEBPB would be unable to gradually increase PPARG expression levels for days since proteins with a 1 hour half-life will have already reached more than 99% of their maximal level within 12 hours of stimulation (Figure 4E, red line). However, a slow-degrading regulator of PPARG would be able to slowly buildup up and increase PPARG levels for days for continuous stimuli (Figure 4E, blue line).

The existence of a slow-degrading PPARG feedback partner would also explain another observed phenomenon: that the adipogenic differentiation system responds very differently to cversus continuous stimuli (Figures 1C-E, 3B-C, 4A-B), filtering out stimuli in the first case and regulating increasing fractions of the cell population to differentiate in the second case. A system with only fast-degrading regulators would only have one response since it would rise rapidly to the same expression level for both oscillating or continuous stimuli and would either trigger differentiation or not for both types of stimuli without a delay (Figure 4F, top). In contrast, a slow-degrading PPARG regulator could rise to different steady-state amplitudes in response to different input pulse durations and thus would be able to keep its steady-state level below a threshold for oscillating stimuli while also being able to rise above the threshold and trigger differentiation for continuous stimulation (Figure 4F, bottom, solid red and blue lines). In other words, the existence of a slow-degrading PPARG regulator would allow a system to selectively prevent differentiation for oscillatory pulsatile stimulation as long as the trough between pulses is sufficiently long (~12 hours) which allows for sufficient degradation of the slow regulator between pulses. The dashed lines in Figure 4F shows how steady-state levels increase for daily pulses of 18-hour and become lower for daily pulses of 6-hour duration. The existence of slow-degrading regulators in the adipogenic transcriptional architecture would enable adipocyte precursor cells to convert daily hormone stimuli of different pulse durations into different longterm steady-state amplitudes of expression of the regulator above and below a threshold. In summary, a slow feedback regulator of PPARG is needed both to increase PPARG levels slowly to the threshold to differentiate (Figures 3B-C, 4E) and to allow PPARG levels to rise to different steady-state levels for oscillating versus continuous stimuli (Figure 4F). Since a transcriptional positive feedback loop between PPARG and CEBPA has been shown to be particularly critical for differentiation (El-jack et al., 1999; Wu et al., 1999) (Figure 2A), an obvious candidate for such a slow regulator is CEBPA. However, CEBPA also has short mRNA and protein half-lives of approximately 1 and 3 hours, respectively, and cannot be the required slow co-regulator (Figure 4G).

### Identification of a slow feedback regulator of PPARG that can mediate the slow buildup in PPARG levels to the threshold to differentiate

Since their mRNA and protein all degrade quickly even in the presence of continuous stimuli, the three core regulators CEBPA, CEBPB, and PPARG (Figure 2A) cannot be alone responsible for the observed slow buildup of PPARG. A plausible alternative mechanism to explain the delayed PPARG increase would be if additional positive feedback regulators of PPARG had a much longer life time. Several such positive regulators of PPARG have been identified (Ahrends et al., 2014; Wakabayashi et al., 2009). At least two have been shown to have long protein lifetimes, FLNA and CEPBZ (Schwanhäusser et al., 2011), and one has been shown to have a long mRNA lifetime FABP4 (Sharova et al., 2009; Spangenberg et al., 2013). To understand how a slow regulator might work in a differentiation system, we focused on FABP4 since previous work showed that it had a particularly strong effect on differentiation of OP9 cells (Ahrends et al., 2014).

We first confirmed that FABP4 mRNA has a long half-life of between 14 to 34 hours (Figure 5A and Figure S6) and that the increase in PPARG level is closely correlated with an increase in FABP4 in individual cells (Figure 5B). Previous studies had established FABP4 is a downstream target of PPARG (Hotamisligil and Bernlohr, 2015) that can positively regulate PPARG expression and adipogenesis (Ahrends et al., 2014; Ayers et al., 2007; Boss et al., 2015). Indeed, siRNA-meditated knockdown of FABP4 suppressed adipogenesis (Figure 5C), and overexpression of FABP4 increased adipogenesis in the absence of adipogenic stimulus in OP9 cells (Figure 5D). Furthermore, identical to the effect of CEBPA knockdown, siRNA-mediated depletion of FABP4 and overexpressing FABP4 resulted in a slowing-down (Figure 5E) and acceleration (Figure 5F) of the normally-observed PPARG increase in response to DMI. Together with previous studies by other groups (Ayers et al., 2007; Boss et al., 2015; Tan et al., 2002a), our results support that FABP4 is in a positive-feedback relationship with PPARG in OP9 cells. Furthermore, the timecourses in Figures 5E and 5F support that slow-degrading FABP4 can mediate the slow buildup in PPARG levels. While there are likely additional slow regulators of PPARG expression, our study argues that FABP4 has a critical role as a slow positive feedback partner that mediates the slow PPARG dynamics in response to oscillatory versus continuous stimuli.

**Figure 5.**
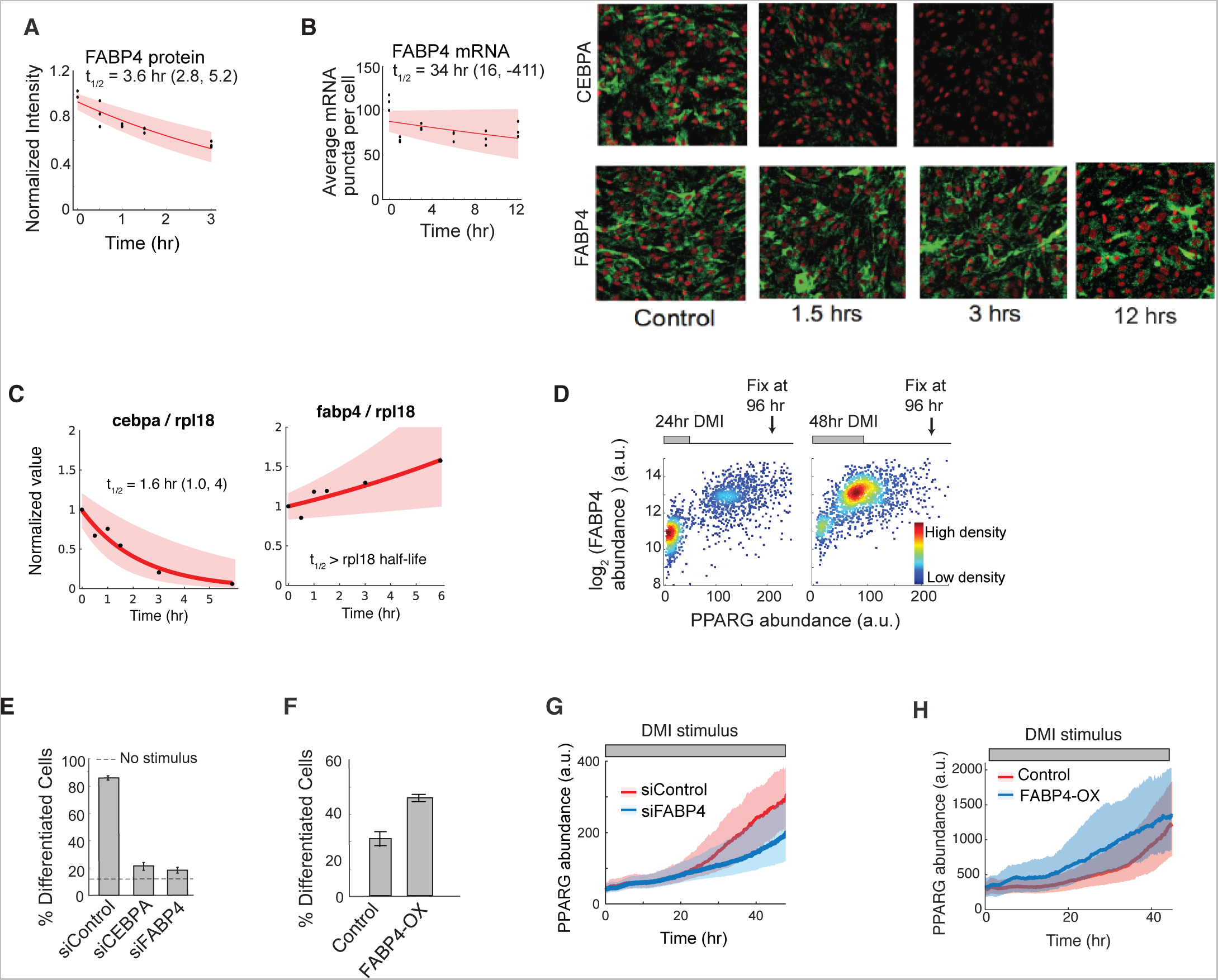
FABP4 is an example of a slow-degrading PPARG regulator that can mediate a slow increase in PPARG expression during adipogenesis. (A) Lifetime analysis of FABP4 protein as in Figure 2G. (B) Lifetime of FABP4 mRNA obtained as in Figure 2G by mRNA FISH analysis of OP9 cells fixed at different times after the addition of actinomycin D. Cebpa mRNA FISH images are shown for comparison. See also Figure S5. (C) Lifetime of FABP4 versus RPL18 mRNA obtained by RT-PCR analysis of OP9 cells collected at different times after the addition of actinomycin D. The lifetime of CEBPA versus RPL18 mRNA is shown for comparison. See also Figure S5. (E) Single cell immunofluorescence analysis showing the mean nuclear PPARG and the mean cytosolic FABP4 values in each cell stimulated with DMI for 24 hours or 48 hours and fixed at 96 hours. Each scatter plot represents approximately 5000 cells and is colored based on cell density. (F) OP9 cells were transfected with siRNA targeting FABP4, CEBPA, or control (YFP), induced to differentiate 24 hours later by usingthe standard protocol of 48-hours of DMI addition followed by 48 hours in media with just insulin. Differentiation was assessed at 96 hours as in Figure 1B. (G) Overexpression of FABP4 increases differentiation even without DMI stimulation. OP9 cells were transfected with constructs expressing either just mcherry(control) or mcherry-FABP4 and left in normal growth media for 5 days. No adipogenic stimulus was added. mcherry is a version of the red fluorescent protein (Shaner et al., 2004). (E-F) Percent of differentiated cells was assesed as in Figure 1B. Barplots show mean +/- s.e.m. from 3 technical replicates, representative of 3 independent experiments. (G-H) Citrine-PPARG cells were transfected in (G) with FABP4-targeted or control(YFP) siRNAs or in (H) with mcherry(control) or mcherry-FABP4. Twenty-four hours after transfection, the cells were induced to differentiate by applying a 48-hour DMI stimulus, and live-cell imaging was carried out to monitor PPARG levels in individual cells. Plotted lines are population median traces with shaded regions representing 25th and 75th percentiles of approximately 700 cells per condition.

### Model calculations show that transcriptional circuits with slow and fast positive feedback can filter periodic stimuli and regulate fractional differentiation

Our results so far suggest that the ability to both reject single and repetitive pulses of stimuli and to regulate increasing fractions of cells in the population to differentiate for continuous stimuli of increasing durations requires a regulatory circuit with fast and slow positive feedback (Figure 6A). To validate this requirement, we carried out simulations using an ordinary differentiation model to predict abundance changes of PPARG driven by the action of combined fast and slow positive feedback (see Methods). In our model, the time constants of the fast and slow feedbacks were set to 3 and 34 hours, respectively, which corresponds to the typical values found for the CEBPA and FABP4 feedbacks to PPARG (Figures 4G and 5A). For a 48-hour continuously-applied stimulus, model calculations showed that a slow feedback partner would mediate a delayed buildup of PPARG past the threshold where differentiation is triggered (Figure 6B). For an oscillating stimulus, the model shows that the concentrations of a slow feedback partner would level off to a much lower steady-state than that reached by continuous stimulus, thereby preventing PPARG from reaching the threshold for differentiation (Figure 6C). It should be noted that for the transcriptional circuit depicted in Figure 6A to be able to filter out circadian input pulses. the slow feedback partner only needs to have a lifetime longer than 12 hours while the fast feedbacks should have a lifetime much shorter than 12 hours.

**Figure 6.**
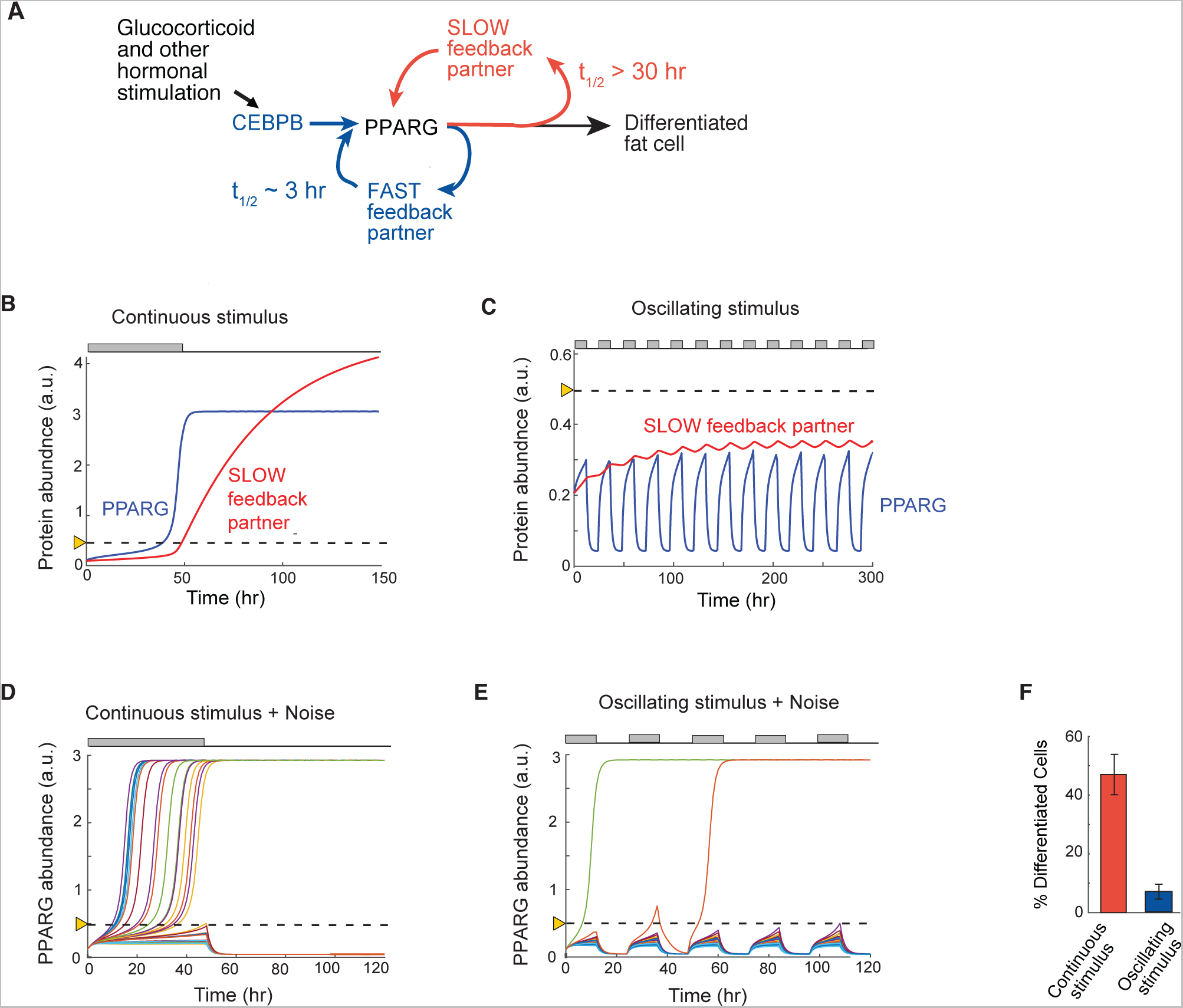
A signaling circuit with a fast and slow positive feedback can trigger differentiation for continuous stimuli and reject daily oscillations. (A) Schematic of the dual feedback loop circuit that rejects circadian inputs but locks cells in the differentiated state for continuous stimuli. (B) Quantitative simulations when a continuous 48-hour stimulus is applied show that the slow positive feedback regulator mediates a slow buildup of PPARG to the threshold for differentiation. The threshold is marked with the dotted line. (C) Same model simulations as in (B) but with an input stimulus applied in circadian 12-hour on / 12-hour off oscillations. The slow regulator builds to a steady-state below the threshold in cells stimulated with circadian hormonal oscillations which keeps levels of its feedback partner PPARG permanently low and below the PPARG threshold where differentiation is triggered. (D and E) Addition of cell-to-cell variability (30% lognormal noise) to a model for a continuous 48-hour stimulus results in high rates of differentiation, but now with variable delays (D). In contrast, addition of cell-to-cell variability to the model with daily oscillating stimuli now allows PPARG levels in a few cells to reach the threshold and result in differentiation. (F) Barplot summarizing 400 simulations with continuous stimulus was applied for 48 hours and 400 stimulations when an oscillating 12-hr on / 12-hr off stimulus was applied. Cells were categorized as differentiated if the PPARG level in that cell rose above the threshold and high PPARG was maintained even when input stimulus was removed. Model results are similar to results from the fixed and livecell experiments shown in Figure 1C-D, 3B, and Figure 4A-B (B-E) Each trace represents a different simulated cell.

The model so far could explain the experimentally-observed rejection of single and repetitive pulses of differentiation stimuli. However, our experiments showed that still a few cells could differentiate for pulsatile stimuli (Figure 1C, 4A). In addition, pulse durations of longer than approximately 12 hours resulted in an increasing fraction of the cell population differentiating (Figure 2B) which also cannot be explained by the above model. We next added noise to our model in order to determine whether cell-to-cell variation (noise) in the slow and fast positive feedback circuit is sufficient to explain why stimulated cells cross the threshold at different times and why periodic stimuli generate low differentiation rates and not zero differentiation. The rationale for the addition of the noise is based on our finding that cell-to-cell variation in PPARG expression enables control of low rates of adipogenesis in a population of precursor cells (Ahrends et al., 2014; Park et al., 2012). Our experimental data in Figures 3B and 3C had shown that cells differentiated at different times when a continuous stimulus was applied. Indeed, adding lognormal stochastic variation to each simulation resulted in individual cells reaching the differentiation threshold with variable delays for continuous stimulation (Figure 6D), reproducing our experimental data in Figure 3B and 3C. Adding the same stochastic noise to the simulations in which differentiation was induced by daily 12-hour on/12-hour off pulses of stimuli showed that cells differentiated only rarely (Figure 6E), again reproducing our experimental data (Figures 4A). Differentiation outcome statistics of oscillating and continuous simulations is shown in Figure 6F. Thus, a circuit with fast and slow positive feedback, together with stochastic variation in PPARG signaling, is sufficient to explain variable delays in cells reaching the threshold and also recapitulates the low differentiation rates observed experimentally for daily oscillations of DMI stimuli.

### Flattening the circadian glucocorticoid oscillations in mice results in a striking increase in adipogenesis without an increase in food intake

To test how oscillating versus continuous levels of glucocorticoid stimuli affect adipogenesis in vivo, we implanted eight-week old C57/Bl6 male mice with pellets that released Corticosterone (Cort) continuously over 21 days (Scheme in Figure 7A and data in Figure 7B). The Cort dose released per day from the pellet was chosen based on previous studies using Cort pellets in mice (Hodes et al., 2012) such that mean Cort levels would not exceed normal mean physiological levels. Mice implanted with Cort pellets showed significant elevated Cort levels in the nadir of the diurnal pattern (08:00-11:00), and also showed lower peak Cort levels (between 17:00-20:00), effectively flattening the level of circulating glucocorticoids without significantly changing the total amount of circulating glucocorticoids in the mice compared to mice implanted with sham pellets (Figure 7B). Such flattening of circadian glucocorticoid oscillations in the bloodstream by implanting Cort pellets has also been demonstrated in rats (Campbell et al., 2011; Dallman et al., 2000). It should be noted that due to the nocturnal waking of mice, the timing of circadian glucocorticoid oscillations in mice is shifted, peaking at approximately 7PM instead of at approximately 7AM as in humans.

**Figure 7.**
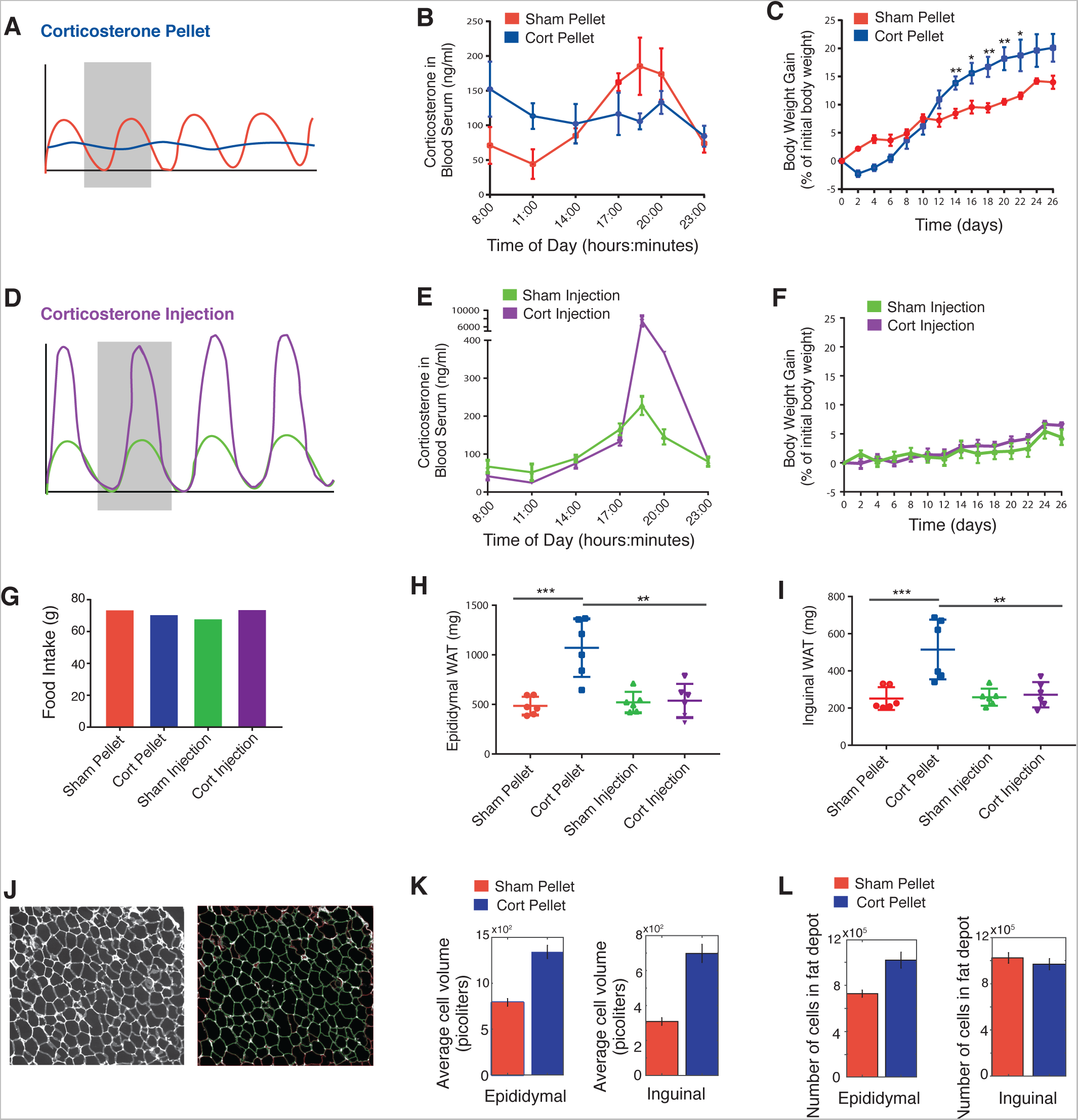
Animals implanted with Corticosterone (Cort) pellets had sustained, flattened elevations in Cort and significant increases in adipose tissue mass. (A) Schematic of pellet implantation protocol to flatten glucocorticoid oscillations in animals. (B) Timecourse of average Cort levels in mice implanted with Cort pellets compared to mice implanted with Sham pellets. N=3, mean +/- s.e.m. (C) Timecourses showing weight of mice implanted with Cort pellets compared to mice implanted with Sham pellets. N=6, mean +/- s.e.m. Unpaired t-test (Sham pellet vs. Cort pellet): * p < 0.05, ** p < 0.01. (D) Schematic of daily injection protocol to selectively increase peak amplitudes of Cort in animals. (E) Peak amplitudes of Cort are greater than 40-fold higher for mice injected daily with Cort compared to mice sham-injected daily with phosphate buffer solution (PBS). N=3, mean +/- s.e.m. (F) Timecourses showing weight of mice injected daily with Cort compared to Sham-injected daily with PBS. N=6, mean +/- s.e.m. (G) Food intake per mouse over the 26-day experimental timecourse for the four specified groups of mice. Each group consisted of 6 mice, housed 3 per cage. Each bar shows the food consumed in the two cages per group divided by 6 mice. (H and I) The weights of both the Epididymal (visceral) and Inguinal (subcutaneous) fat depots were significantly increased for the Cort pellet group but not for the other 3 groups of mice. N=6, mean +/- SD. Unpaired t-test: * p < 0.05, ** p < 0.01, *** p < 0.001. (J) A custom cell contour image analysis program written in MATLAB was used to automatically measure adipocyte size from H & E stained fat tissue slices. Right, example of original H & E image. Left, segmented image showing identified adipocytes in green. Cells touching borders or not meeting shape criteria were excluded. (K-L) Barplots of the average cell volume (K) and number of cells (L) in the Epididymal and Inguinal adipose fat depots in mice treated with Sham or Cort pellets for 21 days. The number of cells were derived by dividing total fat depot weight by the average cell volume. Each bar represents ∼10,000 adipocytes analyzed from 90 images per condition (6 mice with either implanted Sham or Cort pellets × 3 tissue sections per mouse of either Epididymal or Inguinal fat × 5 images per section). Error bars show s.e.m. based on 90 images.

Confirming previous results in which Cort pellets were implanted in rats (Campbell et al., 2011), mice with Cort pellets initially showed a decrease in body weight immediately after receiving the pellets (Figure 7C). However, starting at 2 days after implantation, animals with Cort pellets increased their weight over the next 3 weeks at a faster rate compared to animals receiving the Sham pellets and ended up weighting ~10% more than sham animals on day 26. Sham and Cort groups of implanted animals also had indistinguishable food intake during the experiment (main effect of food: P < 0.05, Figure 7G), arguing that the greater weight increase in Cort animals was likely not a result of increased food intake.

Since our in vitro data had shown that daily pulses with durations less than 12 hours do not result in differentiation independent of their amplitude within a 12-fold amplitude range (Supp. Figure S2C), we tested whether this was the case also in vivo. We generated a 40-fold increase in daily peak amplitude of glucocorticoids compared to Sham-injected mice by injecting Cort into mice at 5PM every day for 21 days (Figures 7D and 7E). Strikingly, despite the greatly increased daily peak levels - and thus total dose - of Cort, we did not observe a significant difference in weight between the Cort‐ and sham-injected animals (Figure 7F), arguing that peak amplitude and total dose of glucocorticoids can change over wide ranges and still not cause an increase in weight as long as the increase in glucocorticoids occurs during a short time window and leaves a sufficiently long time period with low Cort. Again, both the Cort‐ and Sham-injected groups of animals had indistinguishable food intake (Figure 7G). Together, these results suggest that weight gain mostly results from the flattening of glucocorticoid levels rather than the total integrated dose or the peak levels of glucocorticoid activity.

We next tested whether the observed weight gain in mice with flattened glucocorticoid levels is indeed the result of increased fat mass. Markedly, when we dissected animals in the four groups after 26 days, we found that the animals with implanted Cort-pellets - and flattened glucocorticorticoid levels - had significantly greater amounts of both inguinal and epidydimal adipose mass compared to the mice with sham pellets and normal circadian glucocorticoid oscillations (Figures 7H and 7I). Furthermore, despite experiencing daily 40-fold greater peak levels of glucocorticoids for 21 days, the Cort-injected animals showed indistinguishable increases in inguinal and epidydimal adipose mass from sham-injected animals. A summary of the Cort pellet and injection experiments is presented in Table 1.

**Table 1.**
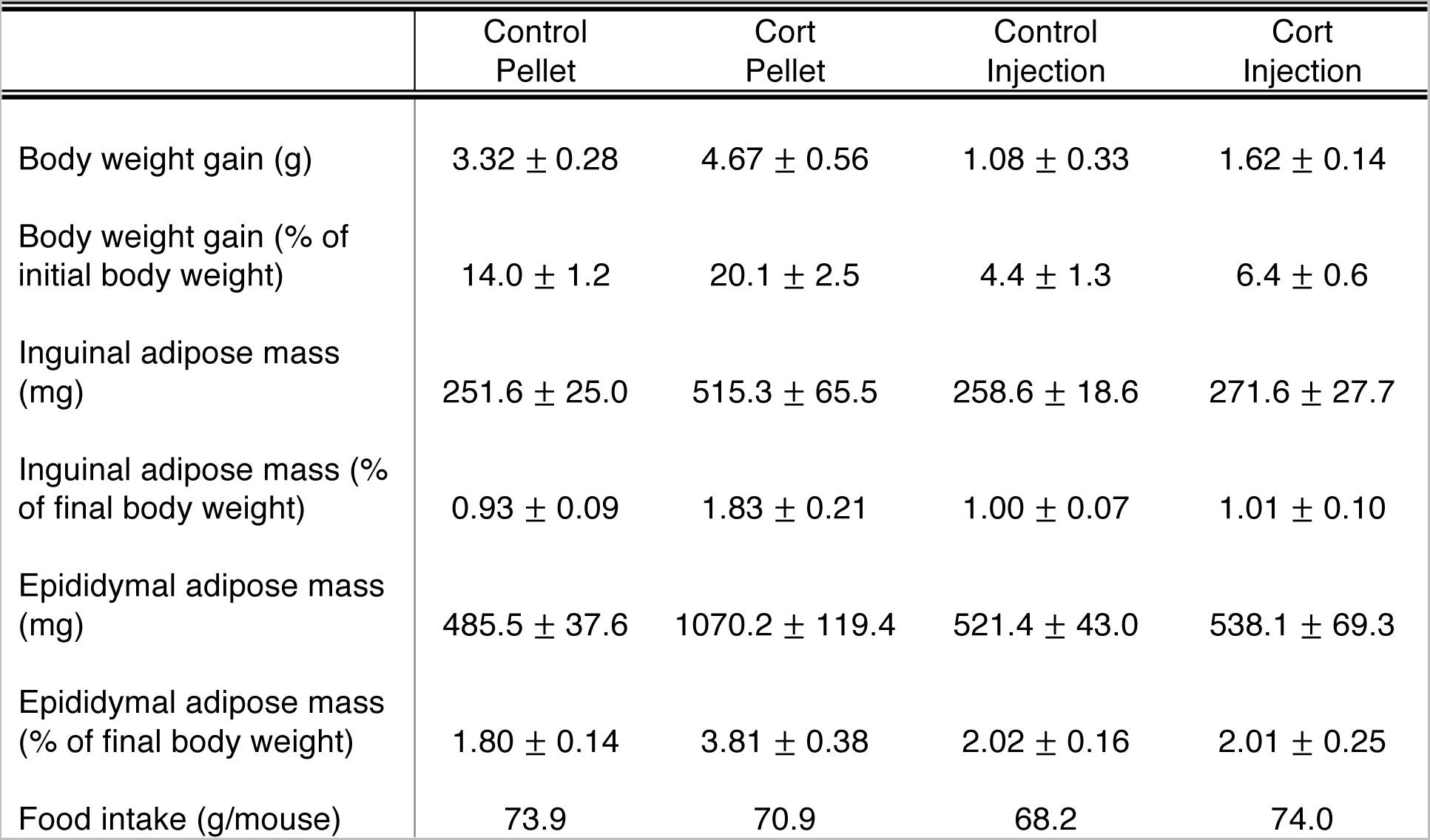
Summary of experiments described in Figure 7 in which circulating Corticosterone (Cort) levels are manipulated in mice using either implanted Cort pellets or daily Cort injections. Values are mean +/- SEM.

To further characterize the adipose tissues in Cort versus Sham-treated mice, we prepared paraffin sections from the inguinal and epididymal fat pads and carried out routine hematoxylin and eosin (H & E) histology staining. We used automated image analysis to compare histology images for the two conditions: Cort-pellet or Sham-pellet mice, and for both types of fat depots: inguinal and epididymal (Figure 7J). We quantified cell volumes since previous studies showed that glucocorticoid-mediated increases in fat pad size are associated with both increased number of cells, as well as with increased cell volume (Campbell et al., 2011; Lee et al., 2014; Rebuffe-Scrive et al., 1992). In both inguinal and epididymal fat pads, animals with implanted with CORT pellets had significantly smaller adipocytes compared with animals implanted with sham pellets (Figure 7K). We then used the cell volume measurement and the respective fat pad weights to derive an estimated number of cells in the fat pads. Epididymal, but not inguinal, fat pads showed a significantly higher number of adipocytes, consistent with previous results in rats (Campbell et al., 2011) (Figure 7L), and suggesting that Cort pellet treatment, which flattens daily glucocorticoid oscillations, results in increased adipogenesis as well as an increase in adipocyte volume.

## DISCUSSION

Our study shows that different preadipocyte model systems (i.e., primary SVF, 3T3-L1, and OP9 cells) fail to differentiate in response to 12-hour on / 12-hour off pulses of hormone stimuli that correspond to the timing of circadian hormone secretions observed in humans and mice. In striking contrast, the same preadipocytes differentiate at gradually increasing rates if the duration of the daily pulses increases from 12 to 18, and 24 hours. Because an increase in pulse duration leads to a shorter trough, or off-period, during each stimulation cycle, our study shows that cells increasingly fail to reset their PPARG level down as pulse durations get longer, thereby increasing the probability that PPARG levels will reach above the threshold and cause cells to continue on the path to the differentiated state. We validate these findings in mice and show that just flattening the circadian glucocorticoid oscillations while keeping the overall circulating glucocorticoid concentrations the same over 21 days caused fat mass to double in the mice compared to mice with normal circadian oscillations. Furthermore, when we raised the peak of the glucocorticoid oscillations 40-fold by injecting corticosterone into mice daily for 21 days, there was no increase in fat mass compared to sham (PBS)-injected mice, providing strong validation that as long as glucocorticoid increases occur within the correct circadian timeperiods, there is only minimal adipogenesis and that it is not increased levels of glucocorticoids, but rather losing the nadirs or "off-periods" that leads to increased fat mass.

We were surprised to find in our live-cell analysis of endogenous CEBPB that the expression of CEBPB increases and decreases along with the daily hormone pulses, reaching each time quickly a maximal level and then dropping again back to basal levels after the hormone stimulus is removed. The closely mirrored relationship between internal CEBPB levels and the external pulse pattern argues that filtering has to occur downstream of CEBPB. We further showed that the life-time of the mRNA and protein of PPARG, CEBPB and CEBPA, are all relatively short lived, and all drop back down rapidly when the input stimulus is removed, which can explain how cells can reset back to their basal state after each of the external hormone pulses as long as the PPARG level in the cell has stayed below the threshold for differentiation. However, these findings did not explain how continuous stimuli can trigger a gradual buildup in PPARG which motivated us to search for slow-acting PPARG regulators. As one such slow-degrading co-factor, we identified FABP4, whose mRNA has a long lifetime of between 14 to 34 hours. We showed that its expression can accelerate PPARG induction and that it is also needed to mediate the slow 24 to 48-hour increase in PPARG levels up to the threshold where differentiation is triggered. This argues for a control principle for differentiation whereby a combined fast and slow positive feedback generates a differentiation system with a delayed and irreversible threshold that can filter out short pulses and circadian hormone stimuli.

FABP4 is one of the most abundant proteins in adipocytes and is a downstream target of PPARG that is highly upregulated during adipogenesis (Hotamisligil and Bernlohr, 2015). Our live-cell analysis of the differentiation decision showed that FABP4 has a critical role in activating PPARG and in controlling the irreversible switch from preadipocyte to adipocyte differentiation. Our results are in line with several other studies that showed that FABP4 upregulates PPARG expression and activity (Adida and Spener, 2006; Ahrends et al., 2014; Ayers et al., 2007; Boss et al., 2015; Tan et al., 2002a). Nevertheless, FABP4’s role in adipogenesis has been controversial since knockout of FABP4 in mice showed no decrease in fat mass (Hotamisligil et al., 1996). However, several studies have since shown that loss of FABP4 in adipocytes can be compensated for by FABP5, a fatty-acid binding protein that is also expressed in adipocytes and has high sequence similarity to FABP4 (Furuhashi et al., 2014; Haunerland and Spener, 2004; Hotamisligil and Bernlohr, 2015; Shaughnessy et al., 2000). In FABP4-knockout mice, FABP5 expression is massively increased: 40-fold at the mRNA level and 13 to 20-fold at the protein level (Coe et al., 1999; Hotamisligil et al., 1996). Indeed when both FABP4 and FABP5 are knocked out in mice, there is a significant reduction in total body adipose tissue mass both on regular and high-fat diet (Maeda et al., 2005).

The transcriptional activity of PPARG is activated by lipids (Tontonoz et al., 1994). FABP4 is one of the most abundant proteins in adipocytes, and several studies have provided evidence that FABP4 has a role in transporting fatty-acid ligands across the cytosol and across the nuclear membrane to PPARG in the nucleus (Adida and Spener, 2006; Ayers et al., 2007; Tan et al., 2002b), thereby enhancing the transcriptional activity of the PPARG. As we point out in the Results section, we are aware that FLNA and CEBPZ are also slow positive feedback regulators, suggesting that FABP4 is likely not the only regulator of differentiation responsible for the filtering.

Since preadipocytes differentiate at low rates in humans and mice (Spalding et al., 2008; Wang et al., 2013) and since a flattening of daily glucocorticoid oscillations leads to increased fat mass and obesity (Campbell et al., 2011; Dallman et al., 2000; Lee et al., 2014), our live-cell measurements provide a molecular basis for strategies to alter the daily timing of glucocorticoid pulses that might be beneficial therapeutically to influence differentiation rates. Specifically, it is suggestive that the dramatic suppression of differentiation for circadian hormone pulse patterns and the much higher rates of differentiation that we found for more sustained hormone signals may provide an explanation as to why conditions such as Cushing’s disease, prolonged stress or long-term glucocorticoid treatments that disrupt normal circadian patterns of hormone oscillations also result in increased obesity. Finally, the molecular filtering mechanism for differentiation we uncovered for adipocytes provides support for the development of temporal therapeutic regimens aimed at changing adipogenic or other hormone pulse durations to control differentiation of precursor cells.

In conclusion, our study introduces a temporal control mechanism for adipogenesis that allows precursor cells to reject normal, daily, oscillating hormone inputs. We show that a dual fast and slow positive feedback system centered on PPARG has the marked characteristic to remain unresponsive to circadian and rhythmic hormone pulses as long as the duration of the trough between pulses remains longer than 12 hours. However, when pulse duration increases and the trough duration becomes shorter, cells convert the duration of the daily pulses into an increasing probability for differentiation until the rate of differentiation becomes maximal for continuous stimulation. Our findings are likely of relevance for many, if not most, differentiation systems since oscillating hormone stimuli are a near universal stimulus pattern in mammalian physiology.

## SUPPLEMENTAL INFORMATION

### AUTHOR CONTRIBUTIONS

Z.B., M.L.Z., S.T., D.H., K.T., and M.N.T. conceived experiments. Z.B., M.L.Z., S.T., D.H., K.T., S.V., and M.N.T. performed experiments, and analyzed data. M.L.Z. and M.C. wrote the image analysis scripts. Z.B., M.L.Z., and M.N.T wrote the paper with input from all authors.

## ACKNOWLEDGEMENTS

This work was supported by National Institutes of Health RO1-DK101743, RO1-DK106241, and P50-GM107615 (to M.N.T.), Stanford BioX Seed Grant funding (to M.N.T.), and T32-NIH T2HG00044 (to M.L.Z.). We thank Sean Collins (UC Davis), James Ferrell, Tobias Meyer, Brian Feldman, Fredric Kraemer, Connie Phong (Stanford University), and members of the Teruel Lab for discussions and critical reading of the manuscript.

## REFERENCES

Adida, A., and Spener, F. (2006). Adipocyte-type fatty acid-binding protein as inter-compartmental shuttle for peroxisome proliferator activated receptor gamma agonists in cultured cell. Biochim. Biophys. Acta 1761, 172–181.

Ahrends, R., Ota, A., Kovary, K.M., Kudo, T., Park, B.O., and Teruel, M.N. (2014). Controlling low rates of cell differentiation through noise and ultrahigh feedback. Science 344, 1384–1389.

Ayers, S.D., Nedrow, K.L., Gillilan, R.E., and Noy, N. (2007). Continuous nucleocytoplasmic shuttling underlies transcriptional activation of PPARgamma by FABP4. Biochemistry 46, 6744–6752.

Bergmann, O., Bhardwaj, R.D., Bernard, S., Zdunek, S., Barnabé-Heider, F., Walsh, S., Zupicich, J., Alkass, K., Buchholz, B.A., Druid, H., et al. (2009). Evidence for cardiomyocyte renewal in humans. Science 324, 98–102.

Boss, M., Kemmerer, M., Brune, B., and Namgaladze, D. (2015). FABP4 inhibition suppresses PPARgamma activity and VLDL-induced foam cell formation in IL-4-polarized human macrophages. Atherosclerosis 240, 424–430.

Campbell, J.E., Peckett, A.J., D’souza, A.M., Hawke, T.J., and Riddell, M.C. (2011). Adipogenic and lipolytic effects of chronic glucocorticoid exposure. Am. J. Physiol. Cell Physiol. 300, C198–209.

Carré, N., and Binart, N. (2014). Prolactin and adipose tissue. Biochimie 97, 16–21.

Chawla, A., Schwarz, E.J., Dimaculangan, D.D., and Lazar, M. (1994). Peroxisome Proliferation-Activated Receptor (PPAR) gamma: Adipose-Predominant Expression And Induction Early In Adipocyte Differentiation. Endocrinology 135, 798–800.

Coe, N.R., Simpson, M. a, and Bernlohr, D. a (1999). Targeted disruption of the adipocyte lipid-binding protein (aP2 protein) gene impairs fat cell lipolysis and increases cellular fatty acid levels. J. Lipid Res. 40, 967–972.

Cristancho, A.G., and Lazar, M.A. (2011). Forming functional fat: a growing understanding of adipocyte differentiation. Nat. Rev. Mol. Cell Biol. 12, 722–734.

Dallman, M.F., Akana, S.F., Bhatnagar, S., Bell, M.E., and Strack, A.M. (2000). Bottomed out: metabolic significance of the circadian trough in glucocorticoid concentrations. Int. J. Obes. 24, S40–S46.

El-jack, A.K., Hamm, J.K., Paul, F., Farmer, S.R., and Pilch, P.F. (1999). Reconstitution of Insulin-sensitive Glucose Transport in Fibroblasts Requires Expression of Both PPAR γ and C / EBP α Reconstitution of Insulin-sensitive Glucose Transport in Fibroblasts. J. Biol. Chem. 274, 7946–7951.

Farmer, S.R. (2006). Transcriptional control of adipocyte formation. Cell Metab. 4, 263–273.

Furuhashi, M., Saitoh, S., Shimamoto, K., and Miura, T. (2014). Fatty Acid-Binding Protein 4 (FABP4): Pathophysiological Insights and Potent Clinical Biomarker of Metabolic and Cardiovascular Diseases. Clin. Med. Insights. Cardiol. 8, 23–33.

Haunerland, N.H., and Spener, F. (2004). Fatty acid-binding proteins - Insights from genetic manipulations. Prog. Lipid Res. 43, 328–349.

Hodes, G.E., Brookshire, B.R., Hill-Smith, T.E., Teegarden, S.L., Berton, O., and Lucki, I. (2012). Strain differences in the effects of chronic corticosterone exposure in the hippocampus. Neuroscience 222, 269–280.

Hotamisligil, G.S., and Bernlohr, D.A. (2015). Metabolic functions of FABPs—mechanisms and therapeutic implications. Nat. Publ. Gr. 11, 592–605.

Hotamisligil, G.S., Johnson, R.S., Distel, R.J., Ellis, R., Papaioannou, V.E., and Spiegelman, B.M. (1996). Uncoupling of Obesity from Insulin Resistance Through a Targeted Mutation in aP2, the Adipocyte Fatty Acid Binding Protein. Science 274, 1377–1379.

Jukam, D., and Desplan, C. (2010). Binary fate decisions in differentiating neurons. Curr. Opin. Neurobiol. 20, 6–13.

Kalmar, T., Lim, C., Hayward, P., Munoz-Descalzo, S., Nichols, J., Garcia-Ojalvo, J., and Arias, A.M. (2009). Regulated fluctuations in Nanog expression mediate cell fate decisions in embryonic stem cells. PLoS Biol. 7, 33–36.

Lee, M.J., Pramyothin, P., Karastergiou, K., and Fried, S.K. (2014). Deconstructing the roles of glucocorticoids in adipose tissue biology and the development of central obesity. Biochim. Biophys. Acta 1842, 473–481.

Lefterova, M.I., Haakonsson, A.K., Lazar, M.A., and Mandrup, S. (2014). PPARgamma and the global map of adipogenesis and beyond. Trends Endocrinol. Metab. 25, 293–302.

Lehmann, J.M., Moore, L.B., Smith-Oliver, T. a, Wilkison, W.O., Willson, T.M., and Kliewer, S. a (1995). An antidiabetic thiazolidinedione is a high affinity ligand for peroxisome proliferator-activated receptor gamma (PPAR gamma). J. Biol. Chem. 270, 12953–12956.

Leliavski, A., Dumbell, R., Ott, V., and Oster, H. (2015). Adrenal clocks and the role of adrenal hormones in the regulation of circadian physiology. J. Biol. Rhythms 30, 20–34.

Loewer, A., and Lahav, G. (2011). We are all individuals: Causes and consequences of non-genetic heterogeneity in mammalian cells. Curr. Opin. Genet. Dev. 21, 753–758.

Luo, X., Ryu, K.W., Kim, D.-S., Nandu, T., Medina, C.J., Gupte, R., Gibson, B.A., Soccio, R.E., Yu, Y., Gupta, R.K., et al. (2017). PARP-1 Controls the Adipogenic Transcriptional Program by PARylating C/EBPβ and Modulating Its Transcriptional Activity. Mol. Cell 65, 260–271.

Maeda, K., Cao, H., Kono, K., Gorgun, C.Z., Furuhashi, M., Uysal, K.T., Cao, Q., Atsumi, G., Malone, H., Krishnan, B., et al. (2005). Adipocyte/macrophage fatty acid binding proteins control integrated metabolic responses in obesity and diabetes. Cell Metab. 1, 107–119.

Mandrup, S., and Lane, M.D. (1997). Regulating adipogenesis. J. Biol. Chem. 272, 5367–5370.

Park, Y.-K., and Ge, K. (2017). Glucocorticoid Receptor Accelerates, but Is Dispensable for, Adipogenesis. Mol. Cell. Biol. 37, e00260–16.

Park, B.O., Ahrends, R., and Teruel, M.N. (2012). Consecutive Positive Feedback Loops Create a Bistable Switch that Controls Preadipocyte-to-Adipocyte Conversion. Cell Rep. 2, 976–990.

Rebuffe-Scrive, M., Walsh, U.A., McEwen, B., and Rodin, J. (1992). Effect of chronic stress and exogenous glucocorticoids on regional fat distribution and metabolism. Physiol. Behav. 52, 583–590.

Rodeheffer, M.S., Birsoy, K., and Friedman, J.M. (2008). Identification of white adipocyte progenitor cells in vivo. Cell 135, 240–249.

Roh, H.C., Tsai, L.T., Lyubetskaya, A., Tenen, D., Kumari, M., Rosen, E.D., Roh, H.C., Tsai, L.T., Lyubetskaya, A., Tenen, D., et al. (2017). Simultaneous Transcriptional and Epigenomic Profiling from Specific Cell Types within Heterogeneous Tissues In Vivo: Cell Reports. CellReports 18, 1048–1061.

Rosen, E.D., and Spiegelman, B.M. (2014). What we talk about when we talk about fat. Cell 156, 20–44.

Rosenfeld, N., Elowitz, M.B., and Alon, U. (2002). Negative autoregulation speeds the response times of transcription networks. J. Mol. Biol. 323, 785–793.

Schwanhäusser, B., Busse, D., Li, N., Dittmar, G., Schuchhardt, J., Wolf, J., Chen, W., and Selbach, M. (2011). Global quantification of mammalian gene expression control. Nature 473, 337–342.

Sharova, L. V, Sharov, A.A., Nedorezov, T., Piao, Y., Shaik, N., and Ko, M.S.H. (2009). Database of mRNA Half-Life of 19977 Genes Obtained by DNA Microarray Analysis of Pluripotent and Differentiating Mouse Embryonic Stem Cells Supplementary data. DNA Res. 16, S1.

Shaughnessy, S., Smith, E.R., Kodukula, S., Storch, J., and Fried, S.K. (2000). Adipocyte metabolism in adipocyte fatty acid binding protein knockout mice (aP2-/-) after short-term high-fat feeding: functional compensation by the keratinocyte [correction of keritinocyte] fatty acid binding protein. Diabetes 49, 904–911.

Siersbæk, R., Nielsen, R., John, S., Sung, M.-H., Baek, S., Loft, A., Hager, G.L., and Mandrup, S. (2011). Extensive chromatin remodelling and establishment of transcription factor “hotspots” during early adipogenesis. EMBO J. 30, 1459–1472.

Spalding, K.L., Arner, E., Westermark, P.O., Bernard, S., Buchholz, B. a, Bergmann, O., Blomqvist, L., Hoffstedt, J., Näslund, E., Britton, T., et al. (2008). Dynamics of fat cell turnover in humans. Nature 453, 783–787.

Spangenberg, L., Shigunov, P., Abud, A.P.R., Cofre, A.R., Stimamiglio, M.A., Kuligovski, C., Zych, J., Schittini, A. V., Costa, A.D.T., Rebelatto, C.K., et al. (2013). Polysome profiling shows extensive posttranscriptional regulation during human adipocyte stem cell differentiation into adipocytes. Stem Cell Res. 11, 902–912.

Spencer, S.L., Gaudet, S., Albeck, J.G., Burke, J.M., and Sorger, P.K. (2009). Non-genetic origins of cell-to-cell variability in TRAIL-induced apoptosis. Nature 459, 428–432.

Tan, N., Shaw, N.S., Vinckenbosch, N., Liu, P., Yasmin, R., Desvergne, B., Wahli, W., and Noy, N. (2002a). Selective Cooperation between Fatty Acid Binding Proteins and Peroxisome Proliferator-Activated Receptors in Regulating Transcription Selective Cooperation between Fatty Acid Binding Proteins and Peroxisome Proliferator-Activated Receptors in Regulating T. Mol. Cell. Biol. 22, 5114–5127.

Tan, N.-S., Shaw, N.S., Vinckenbosch, N., Liu, P., Yasmin, R., Desvergne, B., Wahli, W., and Noy, N. (2002b). Selective Cooperation between Fatty Acid Binding Proteins and Peroxisome Proliferator-Activated Receptors in Regulating Transcription. Mol. Cell. Biol. 22, 5114–5127.

Tchoukalova, Y.D., Sarr, M.G., and Jensen, M.D. (2004). Measuring committed preadipocytes in human adipose tissue from severely obese patients by using adipocyte fatty acid binding protein. Am. J. Physiol. Regul. Integr. Comp. Physiol. 287, R1132–40.

Thompson, N.M., Gill, D.A.S., Davies, R., Loveridge, N., Houston, P.A., Robinson, I.C.A.F., and Wells, T. (2004). Ghrelin and des-octanoyl ghrelin promote adipogenesis directlyin vivo by a mechanism independent of GHS-R1a. Endocrinology 145, 234–242.

Tontonoz, P., and Spiegelman, B.M. (2008). Fat and beyond: the diverse biology of PPARgamma. Annu. Rev. Biochem. 77, 289–312.

Tontonoz, P., Hu, E., and Spiegelman, B.M. (1994). Stimulation of adipogenesis in fibroblasts by PPARgamma2, a lipid-activated transcription factor. Cell 79, 1147–1156.

Van, R.L.R., Bayliss, C.E., and Roncari, D.A.K. (1976). Cytological and enzymological characterization of adult human adipocyte precursors in culture. J. Clin. Invest. 58, 699–704.

Wakabayashi, K., Okamura, M., Tsutsumi, S., Nishikawa, N.S., Tanaka, T., Sakakibara, I., Kitakami, J., Ihara, S., Hashimoto, Y., Hamakubo, T., et al. (2009). The peroxisome proliferator-activated receptor gamma/retinoid X receptor alpha heterodimer targets the histone modification enzyme PR-Set7/Setd8 gene and regulates adipogenesis through a positive feedback loop. Mol. Cell. Biol. 29, 3544–3555.

Wang, L., Walker, B.L., Iannaccone, S., Bhatt, D., Kennedy, P.J., and Tse, W.T. (2009). Bistable switches control memory and plasticity in cellular differentiation. Proc. Natl. Acad. Sci. U. S. A. 106, 6638–6643.

Wang, Q.A., Tao, C., Gupta, R.K., and Scherer, P.E. (2013). Tracking adipogenesis during white adipose tissue development, expansion and regeneration. Nat. Med. 19, 1338–1344.

Weitzman, E.D., Fukushima, D., Nogeire, C., Roffwarg, H., Gallagher, T.F., and Hellman, L. (1971). Twenty-four hour pattern of the episodic secretion of cortisol in normal subjects. J. Clin. Endocrinol. Metab. 33, 14–22.

Windle, R.J., Wood, S.A., Kershaw, Y.M., Lightman, S.L., and Ingram, C.D. (2013). Adaptive changes in basal and stress-induced HPA activity in lactating and post-lactating female rats. Endocrinology 154, 749–761.

Wolins, N.E., Quaynor, B.K., Skinner, J.R., Tzekov, A., Park, C., Choi, K., and Bickel, P.E. (2006). OP9 mouse stromal cells rapidly differentiate into adipocytes: characterization of a useful new model of adipogenesis. J. Lipid Res. 47, 450–460.

Wu, Z., Rosen, E.D., Brun, R., Hauser, S., Adelmant, G., Troy, A.E., Mckeon, C., Darlington, G.J., and Spiegelman, B.M. (1999). Cross-Regulation of C / EBPalpha and PPARgamma Controls the Transcriptional Pathway of Adipogenesis and Insulin Sensitivity. Mol. Cell 3, 151–158.

Yeh, W.C., Cao, Z., Classon, M., and McKnight, S.L. (1995). Cascade regulation of terminal adipocyte differentiation by three members of the C/EBP family of leucine zipper proteins. Genes Dev. 9, 168–181.

